# Functional solubilisation of the β_2_-adrenoceptor (β_2_AR) using Diisobutylene maleic acid (DIBMA)

**DOI:** 10.1101/2020.06.29.171512

**Authors:** C. R. Harwood, D. A. Sykes, B. Hoare, F. M. Heydenreich, R. Uddin, D. R. Poyner, S. J. Briddon, D. B. Veprintsev

## Abstract

The β2-adrenoceptor (β2AR) is a well-established target in asthma and a prototypical GPCR for biophysical studies. Solubilisation of membrane proteins has classically involved the use of detergents. However, the detergent environment differs from the native membrane environment and often destabilises membrane proteins. Use of amphiphilic copolymers is a promising strategy to solubilise membrane proteins within their native lipid environment in the complete absence of detergents. Here we show the isolation of the β_2_AR in the polymer Diisobutylene maleic acid (DIBMA). We demonstrate that β_2_AR remains functional in the DIBMA lipid particle (DIBMALP) and shows improved thermal stability compared to the n-Dodecyl-β-D-Maltopyranoside (DDM) detergent solubilised β_2_AR. This unique method of extracting β_2_AR offers significant advantages over previous methods routinely employed such as the introduction of thermostabilising mutations and the use of detergents, particularly for functional biophysical studies.

## Introduction

G protein coupled receptors (GPCRs) are the largest family of membrane proteins within the human genome and are responsible for modulating a broad range of hormonal, neurological and immune responses. It is well established that GPCRs have a large therapeutic potential. Indeed, GPCRs currently represent 34% of all US food and drug administration (FDA) approved drugs, with 475 drugs targeting over 100 diverse receptors (Hauser, Attwood et al. 2017). The β_2_-adrenoceptor (β_2_AR) is a rhodopsin-like family GPCR (Schioth and Fredriksson 2005) and member of the adrenoceptor family, which signals primarily through coupling the heterotrimeric Gs protein. It is a well-established target for asthma and has become one of the most studied GPCRs with several structural (Wacker, Fenalti et al. 2010, Rasmussen, DeVree et al. 2011, Bang and Choi 2015) and detailed biophysical studies (Manglik, Kim et al. 2015, Gregorio, Masureel et al. 2017) into its activation mechanism.

A prerequisite for completion of biophysical and structural studies is the extraction and isolation of the β_2_AR from its cellular environment. Classically, this has involved the use of detergents, in the case of the β_2_AR and other GPCRs, *n*-dodecyl-β-D-maltopyranoside (DDM) is most often used (Munk, Mutt et al. 2019). However, it is well established that detergent micelles do not recapitulate the environment of the cell membrane and, as such, protein stability is compromised. Moreover, there is strong evidence that phospholipid composition affects β_2_AR function (Dawaliby, Trubbia et al. 2016). Cholesterol in particular appears associated with the β_2_AR in crystal structures (Cherezov, Rosenbaum et al. 2007), and improves β_2_AR stability (Zocher, Zhang et al. 2012) and function (Paila, Jindal et al. 2011). Multiple studies (Leitz, Bayburt et al. 2006)(Whorton, Bokoch et al. 2007) have mimicked the native membrane environment and improved protein stability through reconstitution of membrane proteins in liposomes, amphipols or synthetic nanodiscs, however these all require initial use of detergents to extract the membrane protein from the membrane.

Recently, it was discovered that styrene maleic acid (SMA) copolymer directly incorporates into biological membranes and self-assembles into native nanoparticles, known as Styrene Maleic Acid Lipid Particles (SMALPs) (Knowles, Finka et al. 2009) (Stroud, Hall et al. 2018). This has provided a novel method for the solubilisation of membrane proteins with their native receptor associated phospholipids, whilst avoiding the use of detergents at all stages. SMA has been used to solubilise a range of membrane proteins (Dorr, Koorengevel et al. 2014, Gulati, Jamshad et al. 2014, Sun, Benlekbir et al. 2018) including GPCRs (Bada Juarez, Munoz-Garcia et al. 2020)(Jamshad, Charlton et al. 2015) for both structural and biophysical studies. Such studies either improved protein stability compared to detergent or have allowed extraction of membrane proteins that were previously unstable in detergents. There is, however, evidence that the conformational flexibility of GPCRs within SMALPs is restricted (Mosslehy, Voskoboynikova et al. 2019)(Routledge, Jamshad et al. 2020) therefore differing from the native state of the protein. Furthermore, the high absorbance of SMA copolymer in the far-UV region makes optical spectroscopic studies of membrane proteins that are encapsulated within SMALPs challenging, and SMALPs are known to be disrupted by divalent cations (Gulamhussein, Meah et al. 2019).

An alternative to SMA is Diisobutylene maleic acid (DIBMA), a copolymer which was developed specifically for the extraction of membrane proteins from the intact bilayer (Oluwole, Klingler et al. 2017). Compared to SMALPs, DIBMALPs are believed to have only a mild effect on lipid packing, be compatible with optical spectroscopy in the far UV range, and tolerate low millimolar concentrations of divalent cations (Oluwole, Klingler et al. 2017). This makes DIBMALPs far more amenable for functional biophysical studies. Although the disk size of SMALPs is believed to vary with different ratios of styrene to maleic acid, DIBMALPs are generally thought to have a larger hydrodynamic radius than SMALPs (Oluwole, Danielczak et al. 2017).

In this study we demonstrate isolation of the functional β_2_AR from the mammalian cell membrane using DIBMA, with improved thermal stability compared to conventional detergent based methods.

## Results

### DIBMALP-β_2_AR retains its pharmacology

A time-resolved fluorescence resonance energy transfer (TR-FRET) based ligand binding assay was established to investigate if the β_2_AR remained functional when extracted from the HEK cell membranes into DIBMALPs. Figure **1** shows saturation binding experiments for the fluorescent antagonist *S*-propranolol-red-630/650 (F-propranolol) (Hello Bio, UK) binding membrane-β_2_AR, DDM-β_2_AR and DIBMALP-β_2_AR. The β_2_AR retained ligand binding ability when extracted from the membrane using both DDM and using the copolymer DIBMA. These data showed comparable affinities for F-propranolol binding to the β_2_AR in membranes (pK_d_ = 7.50 ± 0.05), DDM (pK_d_ = 7.10 ± 0.08) and DIBMA pK_d_ = 7.00 ± 0.13), although with slightly reduced affinity in DIBMA compared to membranes (P=0.02, one-way ANOVA and Tukey’s multiple comparison).

**Figure 1:**
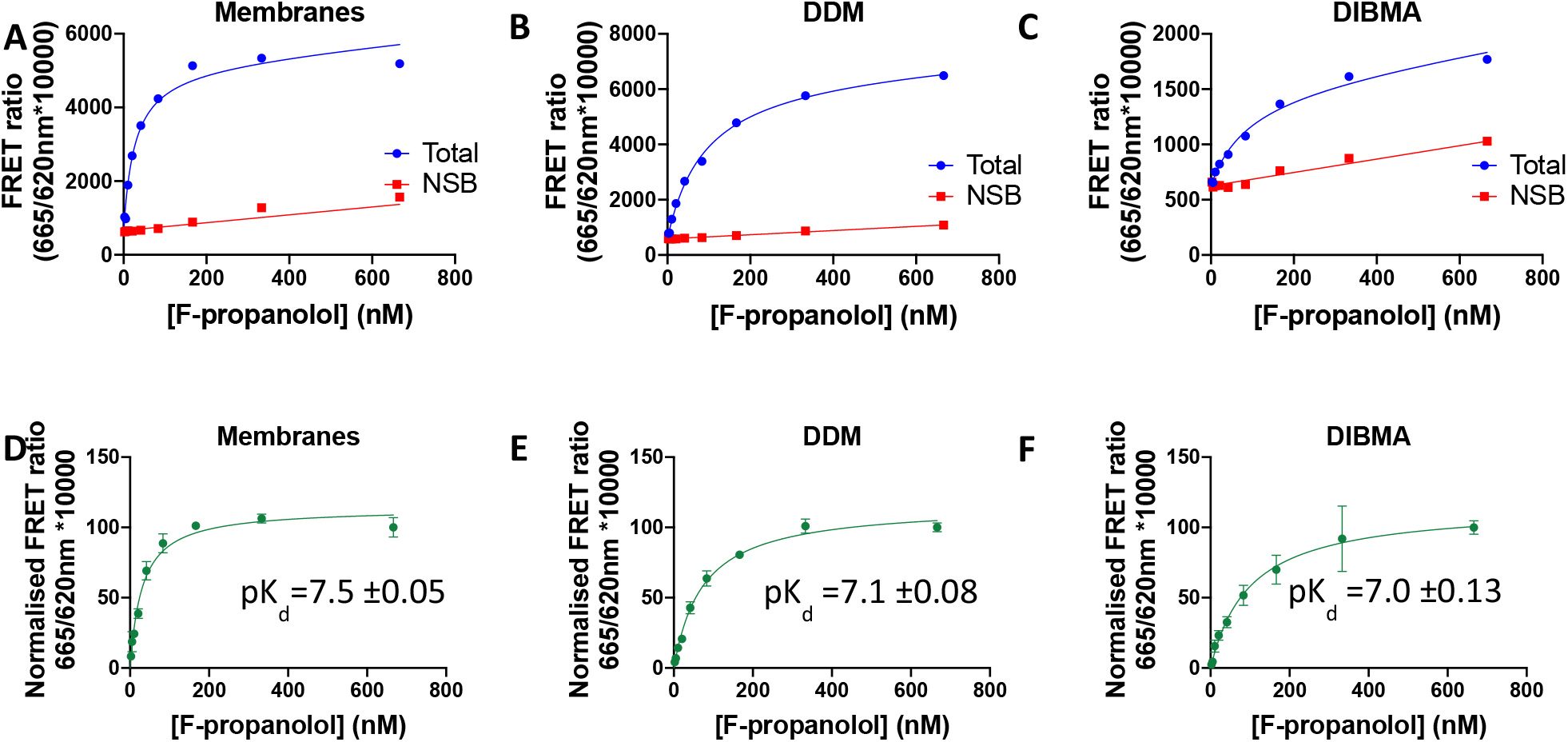
A comparision of F-propranolol binding to β_2_AR in membranes, DDM and DIBMALPs. **A-C)** Representative F-propranolol (2-666nM) equilibrium saturation plots showing total and non-specific binding to the β_2_AR in (A) HEK cell membranes, (B) DDM and (C) DIBMALPs, n=1. **D-F)** Saturation binding curves showing specific binding and associated affinity (pK_d_) values for F-propranolol binding to the β_2_AR in (D) HEK cell membranes, (E) DDM and (F) DIBMALPs, curves show combined normalised data mean ± SEM, n=3.

In order to better understand if the conformational state of the receptor or its ability to adopt different states in DIBMALPs was affected we investigated its pharmacology using the full agonist isoprenaline, the antagonist propranolol and the inverse agonist ICI 118,551 in equilibrium competition binding assays using F-propranolol as the tracer (Figure **2**). Increasing concentrations of each competing ligand produced a reduction in the specific binding of F-propranolol to the β_2_AR in membranes, DDM and DIBMALPs with largely comparable p*K*_i_ values (Table **1**). The only statistically significant difference was between isoprenaline binding to the β_2_AR found in membranes versus the DDM solubilised β_2_AR (p=0.03) (one-way ANOVA and Tukey’s post hoc). The slopes of all curves were similar to 1.

**Figure 2:**
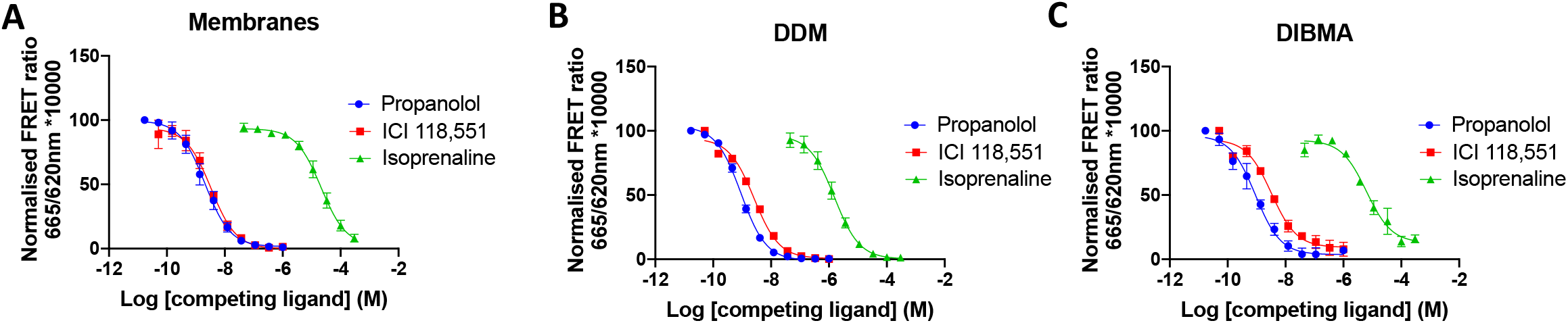
Competition TR-FRET ligand binding studies using F-propranolol as a tracer and unlabelled propranolol, ICI 118,551 and isoprenaline as competitors. **A)** β_2_AR membranes, **B)** DDM-β_2_AR **C)** DIBMALP-β_2_AR, curves show normalized combined data of n=3, error bars show ± SEM.

**Table 1:** A summary of pIC_50_, pK_i_ and Hill slope values for propranolol, ICI 118,551, and isoprenaline obtained through TR-FRET competition binding assays. Values show mean of n=3 individually fitted curves ±SEM.

|  | Membranes |  |  | DDM |  |  | DIBMA |  |  |
| --- | --- | --- | --- | --- | --- | --- | --- | --- | --- |
| | $pIC_{50}$ | $pK_i$ | Slope | $pIC_{50}$ | $pK_i$ | Slope | $pIC_{50}$ | $pK_i$ | Slope |
| **Propranolol** | 8.7 | 9.5 | 1.0 | 9.0 | 9.5 | 1.2 | 9.1 | 9.6 | 0.8 |
| | $\pm 0.13$ | $\pm 0.03$ | $\pm 0.02$ | $\pm 0.04$ | $\pm 0.03$ | $\pm 0.04$ | $\pm 0.10$ | $\pm 0.10$ | $\pm 0.30$ |
| **ICI 118,551** | 8.5 | 9.3 | 1.1 | 8.5 | 8.9 | 1.0 | 8.3 | 9.1 | 1.3 |
| | $\pm 0.10$ | $\pm 0.15$ | $\pm 0.22$ | $\pm 0.02$ | $\pm 0.10$ | $\pm 0.06$ | $\pm 0.15$ | $\pm 0.06$ | $\pm 0.23$ |
| **Isoprenaline** | 4.7 | 5.5 | 1.1 | 5.8 | 6.3 | 1.1 | 5.1 | 5.8 | 1.1 |
| | $\pm 0.12$ | $\pm 0.20$ | $\pm 0.11$ | $\pm 0.06$ | $\pm 0.13$ | $\pm 0.09$ | $\pm 0.18$ | $\pm 0.10$ | $\pm 0.15$ |

### DIBMALP-β_2_AR shows improved stability

Next, we investigated the thermostability of the DIBMALP-β_2_AR using a novel ThermoFRET assay (Figure **3**). Labelling of the SNAP tag on the N terminus of the receptor with Lumi4-Tb allowed thermostability to be investigated without purifying the receptor. β_2_AR unfolding was initially measured by quantifying TR-FRET between Lumi4-Tb and BODIPY™ FL L-Cystine that covalently reacts with cysteines which become exposed as the receptor unfolded (Tippett et al, in preparation).

**Figure 3:**
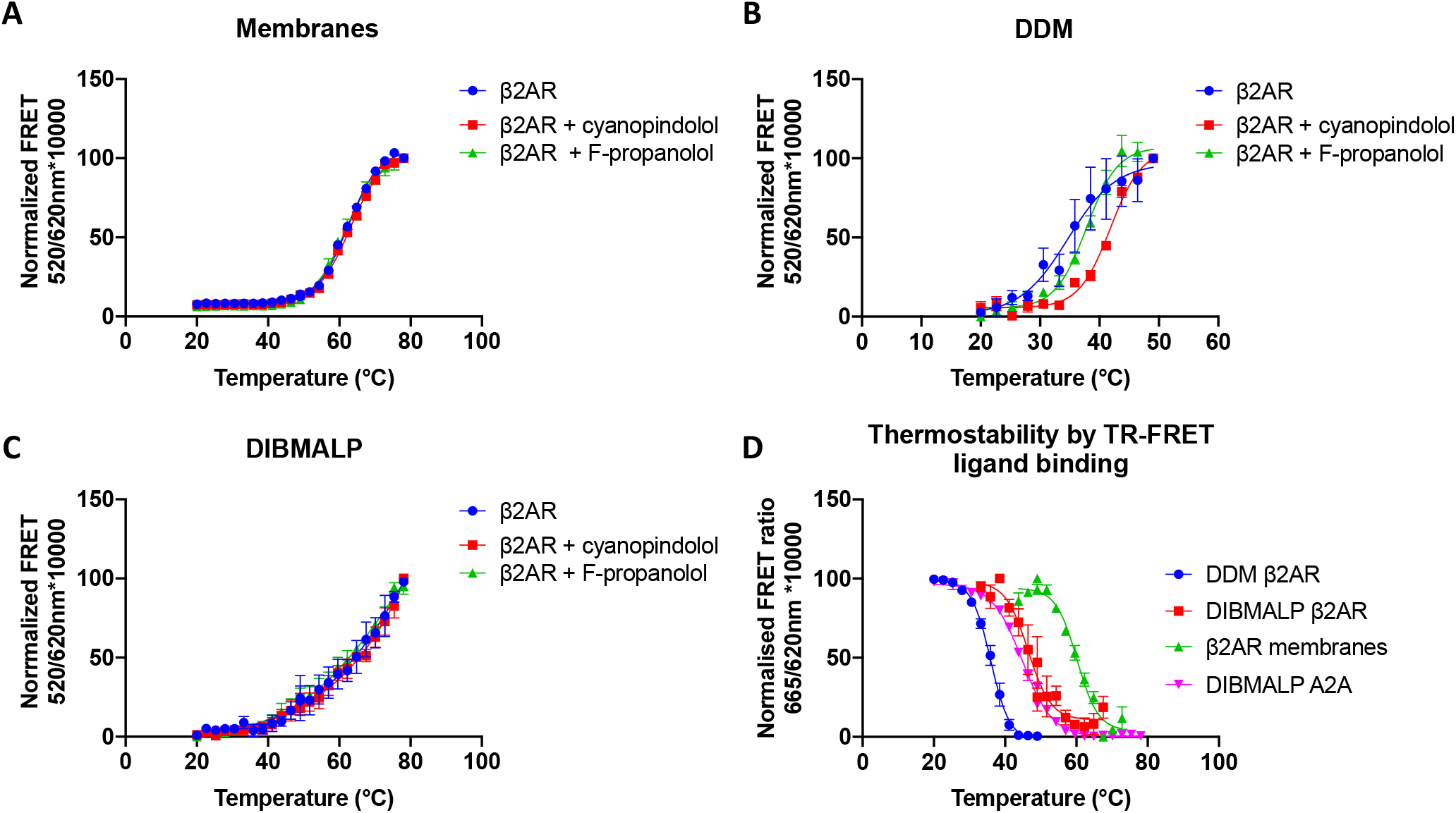
ThermoFRET thermostability curves in **A)** β_2_AR membranes **B)** DDM solubilised β_2_AR **C)** DIBMALP-β_2_AR in the presence and absence of cyanopindolol (100μM) and F-propranolol (200nM). **D)** β_2_AR and A_2_AR TR-FRET thermostability curves obtained by measuring reduction in fluorescent F-propranolol (200nM) and F-XAC (200nM) binding. All curves show normalized combined data, data points show mean ± SEM, for n=3.

**Table 2:**
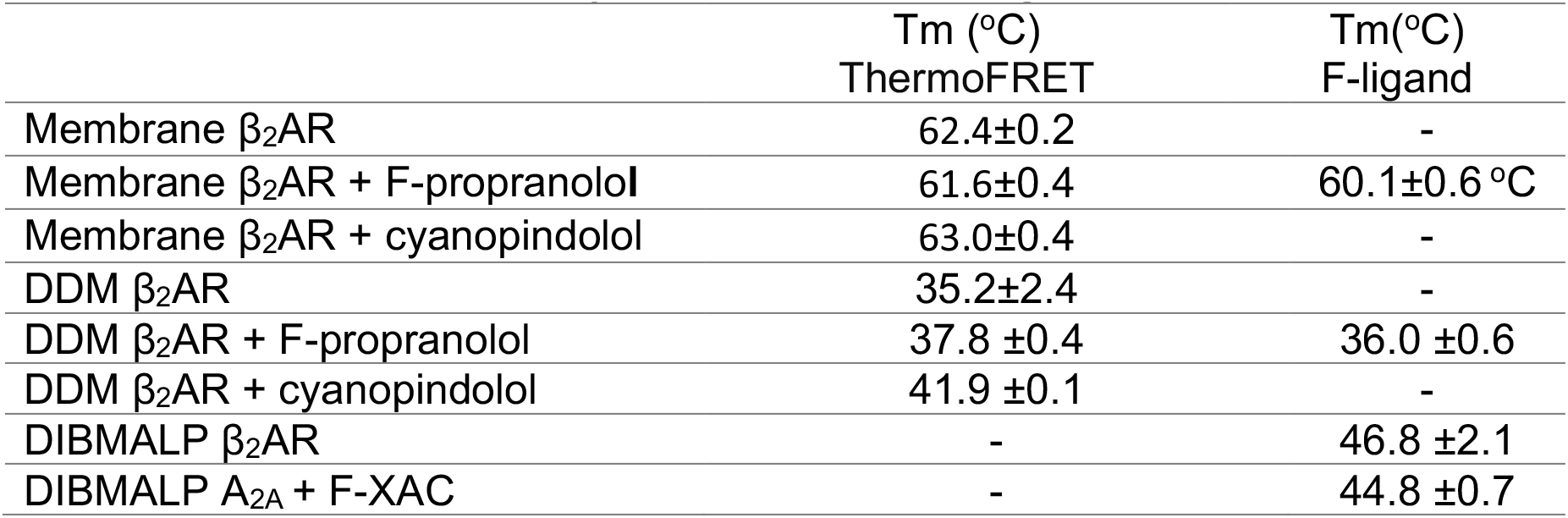
A summary of mean melting temperature (Tm) values ± SEM for β_2_AR in mammalian cell membranes, DDM detergent micelles or DIBMALPs with or without F-propranolol, or cynapindolol measured by HTRF using ThermoFRET or binding of fluorescent ligand F-propranolol as the probe. DIBMALP-A_2A_ thermostability was measured using F-XAC.

|  | T<sub>m</sub> (°C)<br>ThermoFRET | T<sub>m</sub>(°C)<br>F-ligand |
| --- | --- | --- |
| Membrane $\beta_2$AR | 62.4±0.2 | - |
| Membrane $\beta_2$AR + F-propranolol | 61.6±0.4 | 60.1±0.6 °C |
| Membrane $\beta_2$AR + cyanopindolol | 63.0±0.4 | - |
| DDM $\beta_2$AR | 35.2±2.4 | - |
| DDM $\beta_2$AR + F-propranolol | 37.8 ±0.4 | 36.0 ±0.6 |
| DDM $\beta_2$AR + cyanopindolol | 41.9 ±0.1 | - |
| DIBMALP $\beta_2$AR | - | 46.8 ±2.1 |
| DIBMALP A<sub>2A</sub> + F-XAC | - | 44.8 ±0.7 |

Figure **3B** shows the Tm of DDM solubilised β_2_AR as 35.2±2.4 °C. Ligand-induced shifts in thermostability were seen when the DDM solubilised β_2_AR was incubated with F-propranolol (Tm=37.8±0.4°C, p>0.05) and cyanopindolol (41.9 ±0.1 °C, p=0.04) (one-way ANOVA and Tukey’s multiple comparison test). Figure **3A** shows the Tm of β_2_AR in the membrane environment as 62.42±0.2°C. No ligand induced shift was observed when β_2_AR membranes were pre-incubated with F-propranolol or cyanopindolol, this suggests the unfolding of the receptor itself is not directly measurable and perhaps that these data show the disintegration of the membrane itself. Figure **3C** shows TR-FRET thermostability data for the DIBMALP-β_2_AR, this data did not fit a Boltzmann sigmoidal curve as the top end of the temperature range did not plateau. No effect on any part of the curve was observed with the addition of F-propranolol or cyanopindolol. Therefore, as was the case in membranes, the observed thermostability changes in DIBMALPs likely reflect the melting of the lipid particles as opposed to the receptor itself.

We then investigated the thermostability of the β_2_AR by measuring the reduction in TR-FRET binding of F-propranolol over an increasing temperature range (Figure **3D**). The resulting data suggests similar Tm values determined for the membrane-β_2_AR (60.1±0.6 °C) and DDM-β_2_AR (36.0±0.6 °C) to those obtained using BODIPY™ FL L-Cystine in the presence of F-propranolol. Unpaired two-tailed t tests showed no statistically significant differences between membrane-β_2_AR and DDM-β_2_AR Tm values obtained with F-propranolol measured using either TR-FRET method.

Thermostability of DIBMALP-β_2_AR measured by the decrease in F-propranolol binding gave a curve that could be fitted to a Boltzmann with a Tm value of 46.8 ± 2.1 °C. This Tm value is statistically significant from that of membrane-β_2_AR (p=0.0002) and DDM-β_2_AR (p=0.0009) obtained by the same method (one-way ANOVA and Tukey’s multiple comparison test). Therefore, the DIBMALP-β_2_AR shows approximately 10°C improved stability over the conventional DDM-β_2_AR. We also observed differences in the slopes of DIBMALP-β_2_AR and DDM-β_2_AR thermostability curves obtained by this method, these were −3.2 and −2.7 respectively. Additionally, we investigated the thermostability of another rhodopsin-like GPCR, the adenosine 2A receptor (A_2A_R) when solubilised into a DIBMALP using fluorescent adenosine receptor antagonist (F-XAC) (Hello Bio, UK). Measuring the reduction in F-XAC bound to A_2A_R over an increased temperature range gave a Tm value of 44.8°C±0.7, which was not statistically significantly different from that of the DIBMALP-β_2_AR.

## Discussion

The β_2_AR has become the prototypical GPCR for understanding GPCR structure and the molecular basis of signaling (Bang and Choi 2015) (Gregorio, Masureel et al. 2017), these studies have all required the use of detergents to extract the β_2_AR from the plasma membrane. Detergents do not recapitulate the complexity of the native membrane environment and are known to damage membrane proteins. Here, we demonstrate that the polymer DIBMA can be used to extract the β_2_AR from the plasma membrane, together with its native phospholipids, avoiding the use of detergents at any stage.

We used TR-FRET ligand binding studies to show that the β_2_AR remained functional inside the DIBMALP (Figure **2**–**3**). Ligand binding data showed comparable affinity (pK_d_/K_i_) values for the β_2_AR binding F-propranolol, propranolol, ICI 118,551 and isoprenaline solubilised in DIBMA compared to membranes. Although the difference in pK_d_ values for F-propranolol binding membranes-β_2_AR (7.5±0.05) and DIBMALP-β_2_AR (7.0±0.13) was statistically different (P=0.02), this is only a 3 fold difference and the pharmacological importance of this remains to be seen. There was no statistical difference between F-propranolol pK_d_ values for DDM-β_2_AR and DIBMALP-β_2_AR. All ligand binding curves showed one phase binding and a slope of 1 indicating no co-operativity of ligand binding.

While the pK_d_ values for different preparations of the receptor were comparable, the signal amplitude obtained for F-propranolol binding DIBMALP-β_2_AR in TR-FRET experiments was 3-fold lower than for membranes-β_2_AR. This reduction in signal amplitude could be due to an effect of the DIBMA polymer on the TR-FRET, for example fluorescence quenching. Alternatively, it could reflect a lower fraction of the ligand binding capable β_2_AR receptors present compared to the amount of Tb3+ labelled receptor molecules. However, it should be noted that the assay window for DDM-β_2_AR was higher than that of membranes whilst it would be expected that less β_2_AR is functional, suggesting that the solubilization environment can influence the observed signal amplitude. Whilst the concentration of β_2_AR used in each experimental condition was quantified using 620nm emission of Lumi4-Tb, it was not possible to account for difference in Lumi4-Tb quantum yield in the membrane, DDM and DIBMALP environments.

It has been shown that the conformational changes of another class A GPCR, Rhodopsin II in response to activation by light are restricted in SMALPs (Mosslehy, Voskoboynikova et al. 2019). We chose to study the binding of a full agonist (isoprenaline), antagonist (propranolol) and inverse agonist (ICI 118,551) to be able to ascertain if conformational states of the β_2_AR differed in a membrane, DDM micelle, or DIBMALP environment. A substantial increase or decrease in pK_i_ value would demonstrate an increase or decrease in the population of the receptors in the conformational state stabilized by the ligand, and therefore a difference in the conformational landscape of the receptor. As there was no statistically significant difference in pK_i_ values between membrane-β_2_AR and DIBMALP-β_2_AR it can be concluded that the DIBMALP-β_2_AR represents the native conformational landscape of the β_2_AR. The difference in pK_i_ value between DDM-β_2_AR (6.3±0.13) and membrane-β_2_AR (5.5±0.2) for isoprenaline was statistically significant (p=0.03), this may indicate a change in the conformational state of β_2_AR in the DDM micelle compared to its native conformational state. Propranolol, ICI 118,551 and isoprenaline pK_i_ values obtained in this study are in line with the previous studies that investigate the affinity of these compounds for the β_2_AR (Baker 2005) (Sykes, Parry et al. 2014).

Furthermore, we employed a ThermoFRET based thermostability assay to investigate the stability of the DIBMALP-β_2_AR compared to the DDM-β_2_AR. We show the thermostability of DIBMALP-β_2_AR is 10°C higher than that of the DDM-β_2_AR. It was not possible to find any thermostability data for the β_2_AR in synthetic nanodiscs; however, the only other method to show a similar (11 °C) increase in thermostability for β_2_AR is that of thermostabilizing mutations (Serrano-Vega and Tate 2009). Since these mutations also lead to a shift in the β_2_AR’s conformational landscape to the antagonist-bound and inactive form, the DIBMALP-β_2_AR offers a clear advantage for study of receptor function.

Moreover, thermostability data for DIBMALP-β_2_AR using F-propranolol showed a Tm value that was very similar to the Tm of DIBMALP-A_2A_R. In addition, no shift in thermostability was observed for DIBMALP-β_2_AR preincubated with F-propranolol or the high-affinity antagonist cyanopindolol. It therefore seems likely that this Tm value of ~45°C for DIBMALP-β_2_AR corresponds to the melting temperature of the DIBMALP. A Tm of 60.2 ± 0.2°C seen for β_2_AR in membranes was also unaffected by the presence of F-propranolol and cyanopindolol. As this Tm of 60.2°C ±0.2 is statistically significant from that of the DIBMALPs it seems likely that the Tm of ~45°C corresponds to disruption of the protein-lipid-polymer particles, whilst the Tm of 60.2 ±0.2°C corresponds to the melting or disruption of the membrane itself. We also noted a shallower slope for DIBMALP-β_2_AR (−3.2) compared to DDM-β_2_AR (−2.7), this broader transition may reflect the more heterogenous nature of DIBMALPs compared to the detergent micelle.

In summary, here we show the utility of the copolymer DIBMA to solubilise the functional β_2_AR. We show that this method offers improved stability over the use of the conventional detergent DDM and has allowed us to maintain the native environment and pharmacological activity of the β_2_AR.

## Author contributions

**C.R.H** performed the experiments.

**C.R.H.** wrote the manuscript and it was edited by, **D.A.S, D.R.P**, **S.J.B** and **D.B.V**

**D.A.S** gave technical advice and generated reagents.

**R.U** and **D.R.P** gave technical advice.

**B.H and F.M.H.** generated reagents.

**C.R.H, S.J.B and D.B.V** conceived the idea.

**S.J.B and D.B.V** supervised the project

## Acknowledgements

This work was funded by a Medical Research Council (MRC) PhD studentship to C.R.H.

## Supplementary information

### Methods

#### Molecular biology

The construct pcDNA4TO-TwinStrep(TS)-SNAP-β_2_AR was generated by amplification of the SNAP and β_2_AR sequences of the pSNAPf-ADRB2 plasmid (NEB) and insertion into pcDNA4TO-TS using Gibson assembly (Heydenreich, Miljus et al. 2017). The construct pcDNA4TO-TS-SNAP-A_2A_ was generated by amplifying the A_2A_ receptor from the pDNA3.1 SNAP A_2A_ construct described in (Comeo, Kindon et al. 2020) and inserting into pcDNA4TO-TS-SNAP vector using Gibson assembly. This therefore gave the construct pcDNA4TO-TS-SNAP-A_2A_. Both constructs used a signal peptide based on the 5HT_3A_ receptor to increase protein folding and expression.

#### Transfection and mammalian cell culture

pcDNA4TO-TS-SNAP-β_2_AR or pcDNA4TO-TS-SNAP-A_2A_ were stably transfected into T-Rex™-293 cells (Invitrogen) using polyethylenimine (PEI). A mixed population stable line was selected by resistance to 5 μg/mL blasticidin and 20 μg/mL Zeocin. Stable cell lines were maintained in high glucose DMEM (Sigma D6429) with 10% foetal bovine serum (FBS), 5μg/μL blasticidin and 20μg/μL zeocin, at 37°C and 5% CO_2_. When ~70% confluent TS-SNAP-β_2_AR expression was induced with 1μg/mL tetracycline. Cells were left to express for 50hrs before harvesting.

#### Labelling TS-SNAP-β_2_AR with Terbium cryptate

Media was aspirated from T175 flasks and adherent cells washed twice at room temperature with Phosphate Buffered Saline (PBS). Adherent cells were labelled with 100nM SNAP-Lumi4-Tb labelling reagent in Labmed buffer (both Cisbio, UK) for 1 hr at 37°C and 5% CO_2_. Cells were washed twice more with PBS and detached with 5mL non enzymatic cell dissociation solution (Sigma, UK). Cells were pelleted by centrifugation for 10 min at 1000*xg*, supernatant was removed, and cell pellets frozen at −80°C.

#### TS-SNAP-β_2_AR membrane preparation

Cell pellets were thawed on ice and resuspended in 20mL buffer B (10mM HEPES and 10mM EDTA, pH 7.4). Suspensions were homogenised using 6 x 1 sec pulses of a Polytron tissue homogeniser (Werke, Ultra-Turrax). Suspensions were centrifuged at 48,000x*g* and 4°C for 30 min, supernatant was removed and resuspended and centrifuged again as above. Resulting pellets were resuspended in buffer C (10mM HEPES and 0.1mM EDTA, pH 7.4) and frozen at −80°C.

#### Solubilisation of TS-SNAP-β_2_AR using DDM or DIBMA

Membranes were incubated with 3% DIBMA (w/v) (Anatrace, UK) in 20mM HEPES, 10% (v/v) glycerol, and 150mM NaCl, pH 8 at room temperature or 1% DDM (w/v), 20mM HEPES, 5% (v/v) glycerol, and 150mM NaCl, pH 8 at 4°C for 2-3 h. Samples were clarified by ultracentrifugation at 4°C for 1hr at 100,000*xg* for ligand binding assays and 16900*xg* for thermostability assays.

#### TR-FRET ligand binding assays

TR-FRET between the donor Lumi4-Tb and the fluorescent acceptors 633/650 S-propranolol-red (CellAura, UK) (F-propranolol) was measured by exciting at 337nm and quantifying emission at 665nm and 620nm using a PheraStar FSX (BMG Labtech) and HTRF 337 665/620 module (BMG Labtech). Assay buffer consisted of 20mM HEPES, 5% glycerol, 150mM NaCl, and 0.5% Bovine Serum Albumin (BSA), pH8 for DDM solubilised samples 0.1% DDM was used. All binding assays used a final concentration of 1% Dimethyl sulfoxide (DMSO), assay volume of 30μL, 384 well OptiPlates (PerkinElmer, US) and 3μM cyanopindolol was used to determine nonspecific binding (NSB). Receptors were added to plates last and the plates were incubated at room temperature for 45 mins prior to reading. For competition binding assays 100nM of F-propranolol was used for membrane and DDM samples and 200nM F-propranolol for DIBMA samples.

#### ThermoFRET thermostability assays

Solubilised Lumi4-Tb labelled β2AR was incubated with 10μM BODIPY™ FL L-Cystine dye (Molecular Probes, U.S) with or without 200nM F-propranolol or 100μM cyanopindolol, for 15 mins on ice in 20mM HEPES, 150mM NaCl, 5% glycerol, 0.5% BSA, pH8. For DDM samples 0.1% DDM was used. 20μL samples were added to each well of a 96-well PCR plate and incubated for 30 min over a temperature gradient of 20-78°C across the plate using alpha cycler 2 PCR machine (PCRmax, U.K). Samples were transferred to a 384-well proxiplate (PerkinElmer, U.S). TR-FRET between BODIPY™ FL L-Cystine dye and Lumi4-Tb was read by exciting at 337nm and reading emission at 620nm and 520nm using Pherstar FSX and 337 520/620 module (BMG Labtech). F-propranolol and fluorescent XAC (F-XAC) (CellAura, UK) binding was measured using HTRF 337 665/620 module as above.

### Data analysis

#### TR-FRET ligand binding data

Total and NSB for F-propranolol binding to the β_2_AR was fitted to one-site models in GraphPad Prism 8 according to equations 1 and 2.

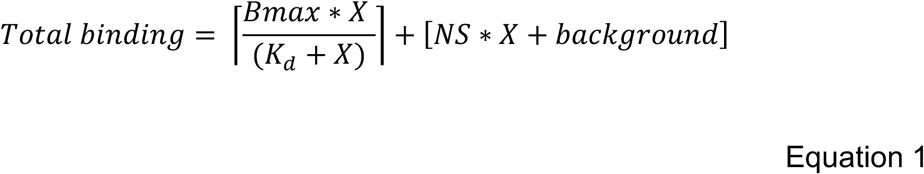

Where:

NS = slope of linear nonspecific binding
Background = Y when X is 0
Bmax = the maximum specific binding
K_d_ = the equilibrium dissociation constant
Y = specific binding
X= concentration of tracer

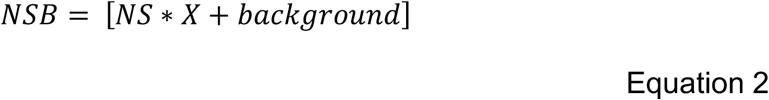

Specific binding of F-propranolol to the β_2_AR was fitted to the one site specific binding model in GraphPad Prism 8 according to equation 3. Final K_d_ values were taken as an average of K_d_ values from individual specific curve fits.

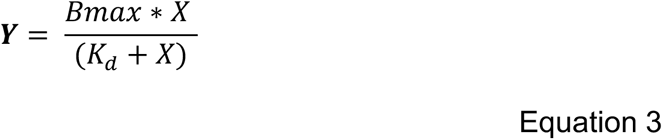

Where:

Y = specific binding
K_d_ = the equilibrium dissociation constant of the labelled ligand

Equilibrium competition binding data was fitted to the One site Fit K_i_ model in GraphPad Prism 8 according to equation 4 and 5. Final K_i_ values were taken as an average of individual curve fits.

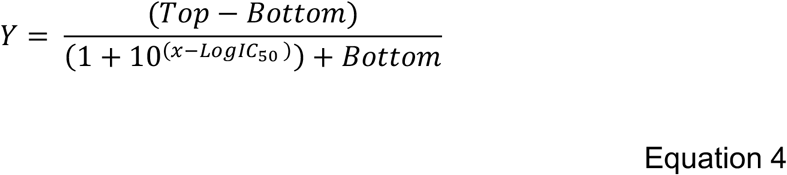

Where:

Y = binding of tracer
IC_50_ = the concentration of competing ligand which displaces 50% of radioligand specific binding.

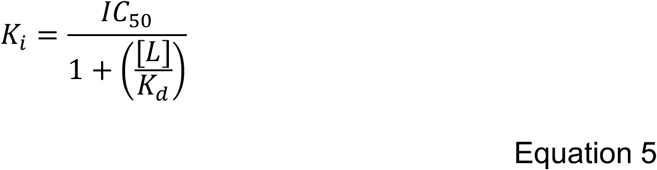

Where:

*K*_i_ = the inhibition constant of the unlabelled ligand
[L] = concentration of labelled ligand
K_d_ = the equilibrium dissociation constant of the labelled ligand.

#### ThermoFRET thermostability curves

All ThermoFRET thermostability data from each experiment was fitted to a Boltzmann sigmoidal curve using GraphPad Prism 8 according to equation 6 to obtain a melting temperature (Tm) value. Final Tm values were taken as an average of Tm values from individual curve fits.

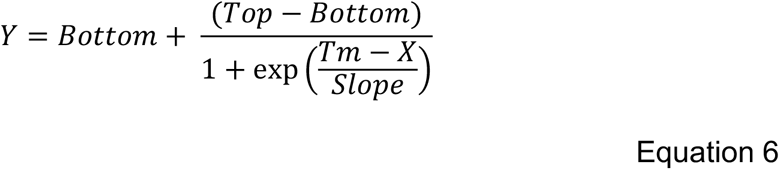

Where:

Y = the relative concentration of proteins in the unfolded state
X = Temperature (°C)
Tm = The temperature at which half the protein of interest is unfolded

### Statistical analysis

Comparison of *T*_m_, *K*_d_ or *K*_i_ values was made using a one-way Analysis Of Variance (ANOVA) test and Tukey’s post hoc multiple comparison test. Statistical comparison of *T*_m_ values obtained with F-propranolol Vs BODIPY™ FL L-Cystine dye was made using an unpaired t test. All statistical analysis was completed in GraphPad Prism 8 and p<0.05 was considered statistically significant.

## Supplementary information

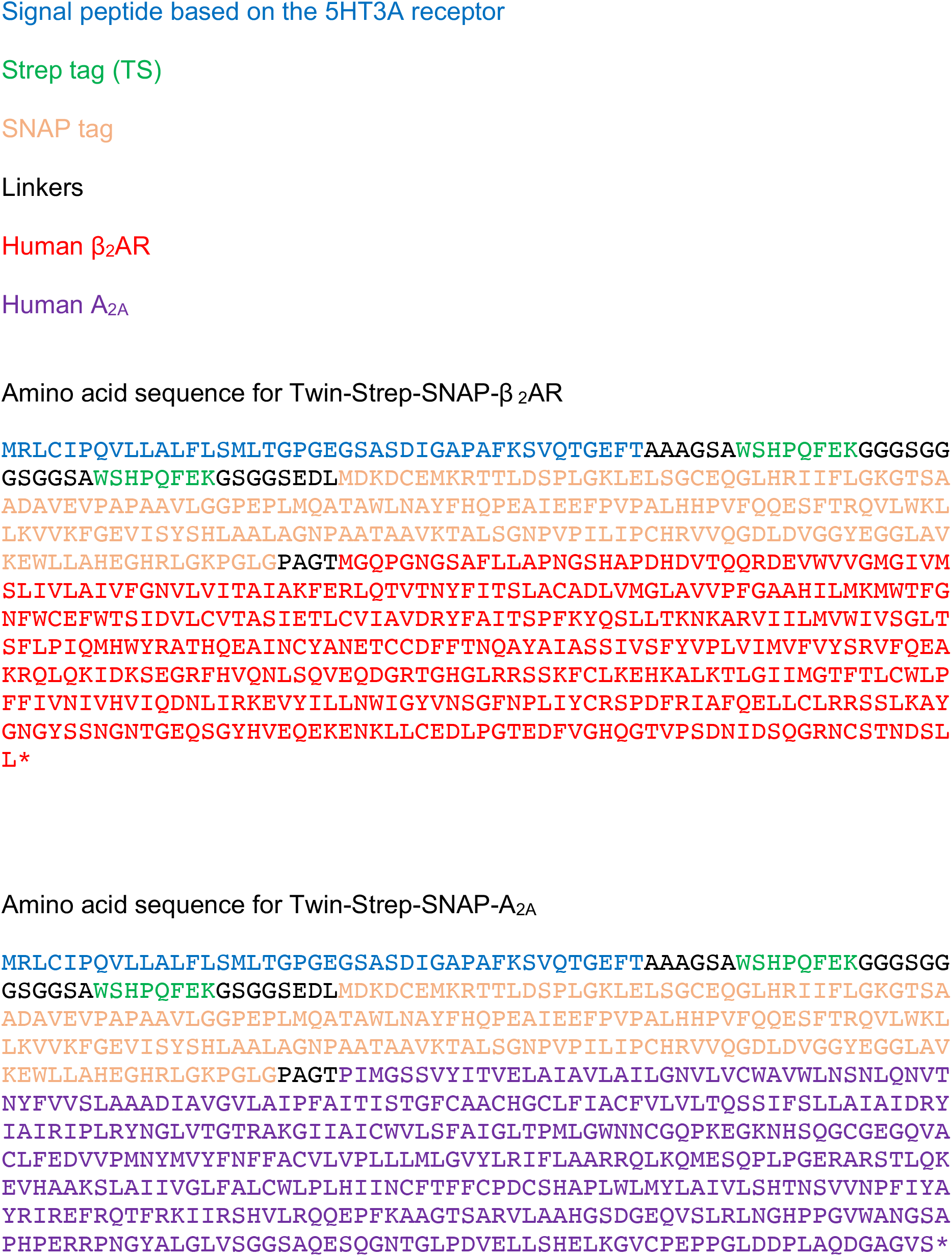
ORF amino acid sequences for Twin-Strep-SNAP-β_2_AR and Twin-Strep-SNAP-A_2A_R

